# Identification of viral dose and administration time in simulated phage therapy occurrences

**DOI:** 10.1101/2022.05.05.490714

**Authors:** Steffen Plunder, Ulrich M. Lauer, Thomas Helling, Sascha Venturelli, Luigi Marongiu

## Abstract

The rise in multidrug-resistant bacteria has sprung a renewed interest in applying phages as antibacterial, a procedure Western practitioners eventually abandoned due to several downfalls, including poor understanding of the dynamics between phages and bacteria. A successful phage therapy needs to account for the loss of infective virions and the multiplication of the hosts. The parameters critical inoculation size (*V_F_*) and failure threshold time (*T_F_*) have been introduced to assure that the viral dose (*v_ϕ_*) and administration time (*t_ϕ_*) would lead to an effective treatment. The problem with the definition of *V_F_* and *T_F_* is that they are non-linear equations with two unknowns; thus, their solution is cumbersome and not unique. The current study used machine learning in the form of a decision tree algorithm to determine ranges for the viral dose and administration times required to achieve an effective phage therapy. Within these ranges, a Pareto optimal solution of a multi-criterial optimization problem (MCOP) provides values leading to effective treatment. The algorithm was tested on a series of microbial consortia that described allochthonous invasions (the outgrowing of a species at high cell density by another species initially present at low concentration) to inhibit the growth of the invading species. The present study also introduced the concept of ‘mediated phage therapy’, where targeting a booster bacteria might decrease the virulence of a pathogen immune to phagial infection. The results demonstrated that the MCOP could provide pairs of *v_ϕ_* and *t_ϕ_* that could effectively wipe out the bacterial target from the considered micro-environment. In summary, the present work introduced a novel method for investigating the phage/bacteria interaction that could help increase the effectiveness of phage therapy.

**Author summary:** Phage therapy is a treatment that can help fight infections with bacteria resistant to antibiotics. However, several phage therapy application have failed, possibly because phages were administered at the wrong time or in insufficient amounts. The present study implemented a machine learning protocol to correctly calculate the administration time and viral load to obtain effective phage therapy. Four simulated microbial consortia, including one case where the pathogen was not directly a phage’s host, were employed to prove the procedure’s concept. The results demonstrated that the procedure is suitable to help the microbiologists to instantiate an effective phage therapy and clear infections.

## Introduction

The rise in multidrug-resistant (MDR) bacteria has sprung a renewed interest in phage therapy, an alternative to the use of antibiotics for the treatment of bacterial infections first introduced by Felix d’Hérelle at the beginning of the 20^th^ century [1]. D’Hérelle isolated phages from environmental sources and used them to treat dysentery and plague, relying on the lytic capability of these viruses towards their bacterial hosts. D’Hérelle’s method was soon adopted by clinical practitioners around the world. Bacteria can develop resistance to their predatory viruses, but the isolation of new strains of phages is much easier to achieve than the production of new types of antibiotics, explaining why there is currently a substantial research interest in phage therapy [2].

Despite being available before the introduction of sulfonamides and antibiotics, the application of phages as antibacterial was abandoned by western practitioners due to several downfalls, including selection of poorly virulent phagial strains, incorrect phage-typing of the infectious bacteria, underestimation of the insurgence of phage-resistance, and presence of bacterial debris in the phagial preparations [3,4]. Furthermore, an inadequate understanding of the dynamics between parasites (phages) and hosts (bacteria) might have led to therapeutic failure [5]. Specifically, such a misinterpretation could have led to erroneous treatment’s dosage and timing.

Any virus requires a minimal number of susceptible hosts to sustain its replication, indicated by the herd immunity threshold 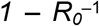, where *R_0_* is the basic reproduction number specific for each parasite (including viruses) [6]. Payne and Jansen at the University of Oxford, UK, contextualized the phagial *R_0_* with the growth rate of the bacterial hosts. In general, phage therapy differs fundamentally from antibiotic therapy because viruses can expand in number. Failing to account for the replicating nature of phages can lead to therapeutic failure. Payne and Jansen described the pharmacokinetic model for phage therapy with the following model [7,8].

The first parameter to consider in phage therapy is the minimal concentration of bacterial host required to sustain a phagial infection. This quantity, known as ‘proliferation threshold’ (*X_P_*), is defined as:

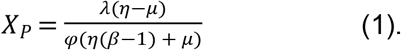

*X_P_* is constant for each pair of phage and bacterial host and it is defined by the life-history traits *μ* (the growth rate of the bacterial host), *φ* (the transmission coefficient of the phage), *λ* (the loss rate of the free phages), *η* (the lysis rate of the infected hosts, that is the reciprocal of the latency time *τ*), and *β* (the burst size of the phage). *X_P_* is reached at a time known as ‘proliferation onset time’ (*T_P_*) defined as:

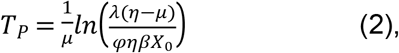

where *X_0_* is the initial concentration of the host. If the phages are administered before *T_P_*, the phages will not replicate and the therapy will be known as ‘passive’. The outcome of the passive therapy depends on the amount of phage inoculated (*v_ϕ_*). If *v_ϕ_* is too low, the therapy will be ineffective because the virions will be removed from the system (either by natural decay of the virions or wash-out caused, for instance, by peristalsis) before the phage/bacterium infection cycle could be established. On the other hand, if *v_ϕ_* is high enough, phages will massively lyse their hosts and the treatment will resemble antibiotic or sulfonamides applications where there is no amplification of the antibacterial agent. To establish passive therapy, *v_ϕ_* must be greater than an ‘inundation threshold’ (*V_I_*), defined as *μ/φ*. Moreover, *v_ϕ_* must also be above a ‘clearance threshold’ (*V_C_*) to effectively wipe out all the bacteria otherwise there will be a proportion of bacteria escaping the phagial infection. *V_C_* is, therefore, a particularly critical quantity and it is defined as:

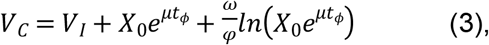

where *t_ϕ_* is the time of inoculation. It is evident from the formula that *V_C_* is greater than *V_I_* due to the terms that account for the expansion of the bacterial population and the loss of virions.

If phages are inoculated after *T_P_*, the phage therapy is defined as ‘active’ because an infection cycle can be sustained and there will be an increase in the number of phages. However, the viral replication is counterbalanced by the loss of virions and the proliferation of the bacteria; therefore, a successful active phage therapy will require an inoculum greater than a ‘critical inoculation size’ (*V_F_*):

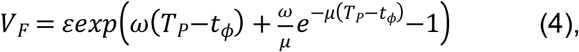

where *ε* is the dilution factor to obtain one phage in the system. Successful active phage therapy is attained if the inoculum is administered after a critical time has been reached, the ‘failure threshold time’ (*T_F_*):

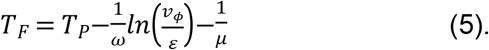

To sum up:

- *t_ϕ_* < *T_P_, v_ϕ_* < *V_I_: passive therapy, ineffective*. There are too few bacteria to establish a sustained infection cycle and the virions are wiped out from the system before the bacterial population is significantly reduced.
- *t_ϕ_* < *T_P_, v_ϕ_* < *V_C_: passive therapy, partially effective*. There are too few bacteria to establish a sustained infection cycle and the virions can reduce the number of bacteria, although no clearance is reached.
- *t_ϕ_* < *T_P_, v_ϕ_* > *V_C_: passive therapy, effective*. There are too few bacteria to establish a sustained infection cycle, but the virions can wipe out the bacterial population.
- *T_F_* < *t_ϕ_* < *T_P_, v_ϕ_* ≤ *V_I_: delayed active therapy, effective*. Even though, initially, there are not enough bacteria to establish a sustained infective cycle, the virions are staying long enough in the environment to lead to active therapy after a delay.
- *t_ϕ_* > *T_P_, v_ϕ_* ≤ *V_I_: active therapy, effective*. There are enough bacteria to establish a sustained infection cycle and, even if a proportion of virions is wiped out from the system, there will be a second wave of phages to maintain the infection cycle until clearance is reached.

The problem with the definition of *V_F_* and *T_F_* is that both *t_ϕ_* and *v_ϕ_* are exactly the sought unknown quantities required to achieve effective therapy. Thus, both Eq. 4 and 5 are non-linear equations with two unknowns; as such, their solution is cumbersome and, without further conditions, not unique.

The aim of the present work was to use a numerical approach to identify *V_F_, T_F_, t_ϕ_*, and *v_ϕ_*. A decision tree algorithm was developed to explore the different outcomes of microbial consortia undergoing phage treatment and to identify the best pairs of *v_ϕ_* and *t_ϕ_* for achieving either active or passive treatment. The identification of a *v_ϕ_/t_ϕ_* pair will facilitate the microbiologist’s work in implementing an effective therapy. The algorithm was tested on a series of microbial consortia: (1) the scenario described by Payne and Jansen in their study on phage therapy; (2 and 3) dual bacteria combinations; (4) two species boosting each other’s fitness.

## Results

In the following sections, decision trees will be built on selected cases describing allochthonous invasions (Fig 1). The trees defined the limits for each type of phage therapy (passive, active, or delayed), providing values equivalent to *T_F_* and *V_F_* (Table 1). Moreover, a pair of viral load and administration time, equivalent to the parameters *v_ϕ_* and *t_ϕ_*, was determined by a multi-criteria optimization problem to provide the user with convenient values for implementing the chosen treatment.

**Fig. 1.**
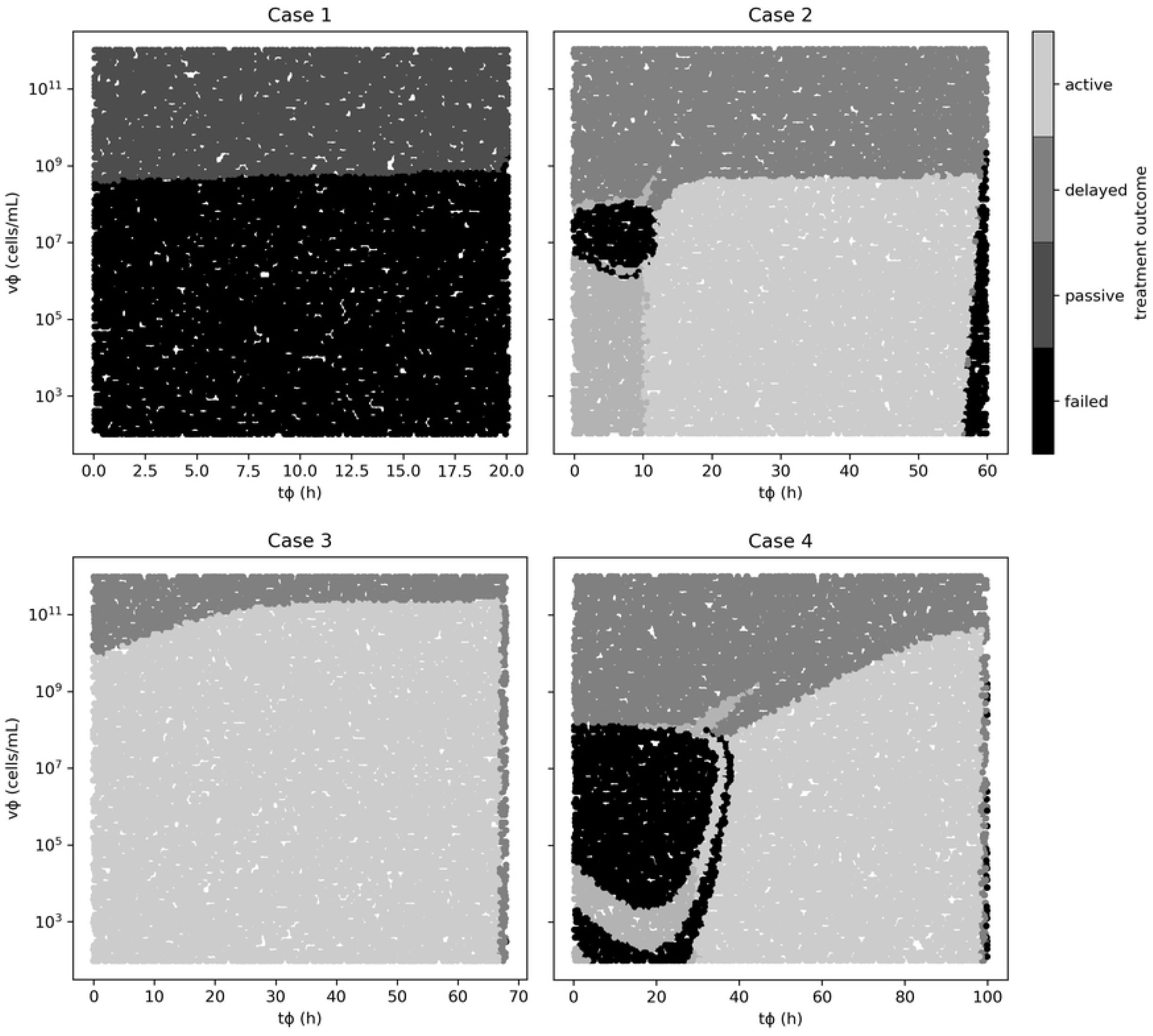
Ensemble simulations for the cases analyzed in the present study. The treatment outcome was evaluated by randomly selecting pairs of viral concentrations and administration times by solving the microbial consortium’s ODE system. Each run pair is color-coded according to the treatment’s outcome. Note that similar outcomes cluster together, and the borders (or margins) of the clusters provide the critical values for the treatments, which are equivalent to *V_F_* and *T_F_*.

**Table 1.** Summary of the phage therapy outcomes obtained by decision tree approach for the cases presented in the present study.

| Case | Microbial consortium | Phage | Outcome (accuracy) | $v_\phi$ range (PFU/mL) | $t_\phi$ range (h) |
| --- | --- | --- | --- | --- | --- |
| 1 | Hypothetical | Hypothetical | Passive (99.5%) | $\geq 5.66 \times 10^8$ | $\geq 0.05$ |
| 2 | *E. coli* vs *P. aeruginosa* | T4 | Active (98.0%) | $\leq 4.50 \times 10^8$ | 11.45-57.05 |
| | | | Delayed (100%) | $\leq 1.42 \times 10^6$ | $\leq 11.45$ |
| | | | Passive (99.3%) | $\geq 4.50 \times 10^8$ | $\leq 59.35$ |
| 3 | *E. coli* vs *A. vinelandii* | T4 | Active (100%) | $\leq 8.97 \times 10^9$ | $\leq 66.75$ |
| | | | Passive | $\geq 1.79 \times 10^9$ | Any |
|  |  |  | (100%) |  |  |
| 4 | *S. mutans*/*C. albicans* vs<br>*L. reuteri* | $\lambda$ | Active<br>(97.3%) | $\leq 1.42 \times 10^8$ | 32.95-98.25 |
| | | | Passive<br>(100%) | $\geq 1.79 \times 10^{10}$ | Any |

### Case 1: hypothetical bacterium and phage

Payne and Jansen described the growth of a hypothetical bacterium and the administration of its phage, highlighting four main treatment outcomes: failed, passive, active, and delayed [5]. In the present study, the failed outcome was used as a base to implement an effective passive therapy. The parameters of the simulation, derived from the Payne and Jansen’s study, were as follows. Initial concentration of bacteria (*X_0_*): 1,000 colonies forming units per milliliter (CFU/mL); phage inoculum (*v_ϕ_*): 10^8^ PFU/mL; time of inoculum (*t_ϕ_*): 2.5 hours; bacterial growth rate (*μ*): 0.500 h^−1^; phagial adsorption rate (*δ*): 10^−7^ min^−1^; reciprocal of latency time (*η*): 5 min; burst size (*β*): 100 PFU; phage decay (*λ*): 5 day^−1^. The bacterial growth was adapted to account for a logistic growth with a carrying capacity (*κ*) of 6.5 × 10^6^ CFU/mL. The outflow rate (*ω*) was set to 0.15 mL/h, and the simulation time-frame was 20 h.

Based on the above parameters, *X_P_* was calculated at 450,000 CFU/mL with a corresponding *T_P_* of 12.2 h. However, *T_P_* did not intersect the growth curve calculated for the hypothetical bacterium but, by regression, the intersection point was at 16.9 h (Fig 2A). The decision tree algorithm developed herein reported only one effective outcome, passive. The margins of this treatment comprised a viral load above 5.66 × 10^8^ PFU/mL with any administration time. The Pareto optimal pair of viral load and administration time was identified as 1.99 × 10^10^ and 3.4 h. The results of the therapy clearly illustrated the characteristics of an effective passive approach: there was no increase in phage density with respect to the initial input and there were no bacteria left in the environment at the end of the simulation, indicating that the infection had been cleared as required (Fig 2B).

**Fig. 2.**
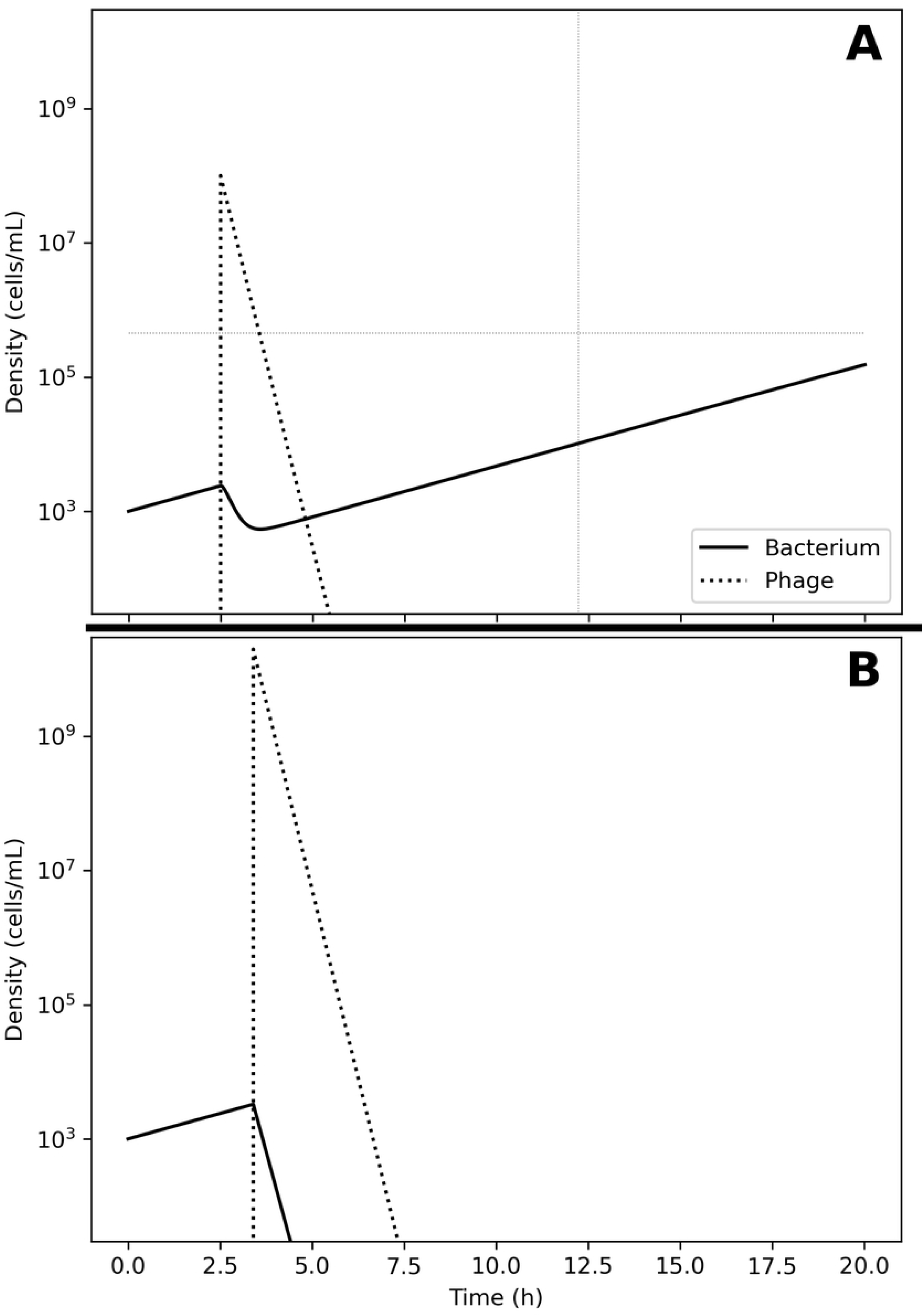
Model of the competition between hypothetical bacteria and phages. Interaction between an ideal bacterium and its phage. A. Failed therapy. The simulation shows a passive therapy, since there was no amplification of the phages over the initial load, In addition the therapy failed because the virions were depleted from the system before the bacterium could be cleared. To note the decrease in bacterial concentration after the application of *v_ϕ_* = 10^8^ phages at *t_ϕ_* = 2.5 h and the increase in density of the escaped bacteria. B. Effective therapy. Implementing the decision tree allows determining the *v_ϕ_* (2.0 × 10^10^) and *t_ϕ_* (3.4 h) values necessary to obtain a successful passive therapy.

A dynamic plot was implemented to actively explore the role of the different parameters in modeling phage therapy (Suppl. File 1). The figure shows that the outcome of the phagial administration is strongly dependent on the parameters used in the computation, highlighting the fact that phage therapy is case-specific.

### Case 2: Escherichia coli vs Pseudomonas aeruginosa

The growth of *Escherichia coli* C-8 and *Pseudomonas aeruginosa* PAO283 was described by Hansen and Hubbell in 1980 using batch cultures [9]. The life-history traits reported by this study for these bacteria were as follows. *E. coli:* yield (*Y*) 2.5 × 10^10^ cells per gram (cell/g) of limiting substance; half saturation constant (*K_S_*) 3.0 × 10 ^−6^ grams per liter (g/L) of limiting substance; *μ_max_* = 0.810 h^−1^. *P. aeruginosa: Y* = 3.8 × 10^10^ cell/g; *K_S_* = 3.0 × 10^−6^ g/L; *ν_max_* = 0.910 h^−1^. The bacteria were growth in 100 mL flasks containing minimal medium with tryptophan as limiting nutrient, provided at an initial concentration of 1.0 × 10^−4^ g/L. The growth rates were calculated according to Eq. 11 (*Materials and Methods* section): *μ* = 0.790 for *E. coli* and *ν* = 0.220 for *P. aeruginosa*. The carrying capacity *κ* was estimated from the original graph at 6.5 × 10^6^ cells/mL. The initial seed of bacteria was extracted from the original graphs: *E. coli*, 334 cells/mL; *P. aeruginosa*, 88,516 cells/mL. These quantities gave a *P. aeruginosa/E. coli* ratio of 265, in line with the reported 200 to 1 for the initial densities of these bacteria. *Escherichia coli* outgrew *P. aeruginosa* about 9.2 hours after the beginning of the experiment and the latter was wiped out in about 60 hours. *X_P_* was calculated to 10,556 cells and *T_P_* at 4.4 hours after the beginning of the experiment. As in case 1, the intersection between *X_P_* and the growth curve of *E. coli* did not occur at *T_P_;* by regression, the intersection point was calculated at 5.5 h, that is only 1.1 hours after *T_P_* (Fig 3A).

**Fig. 3.**
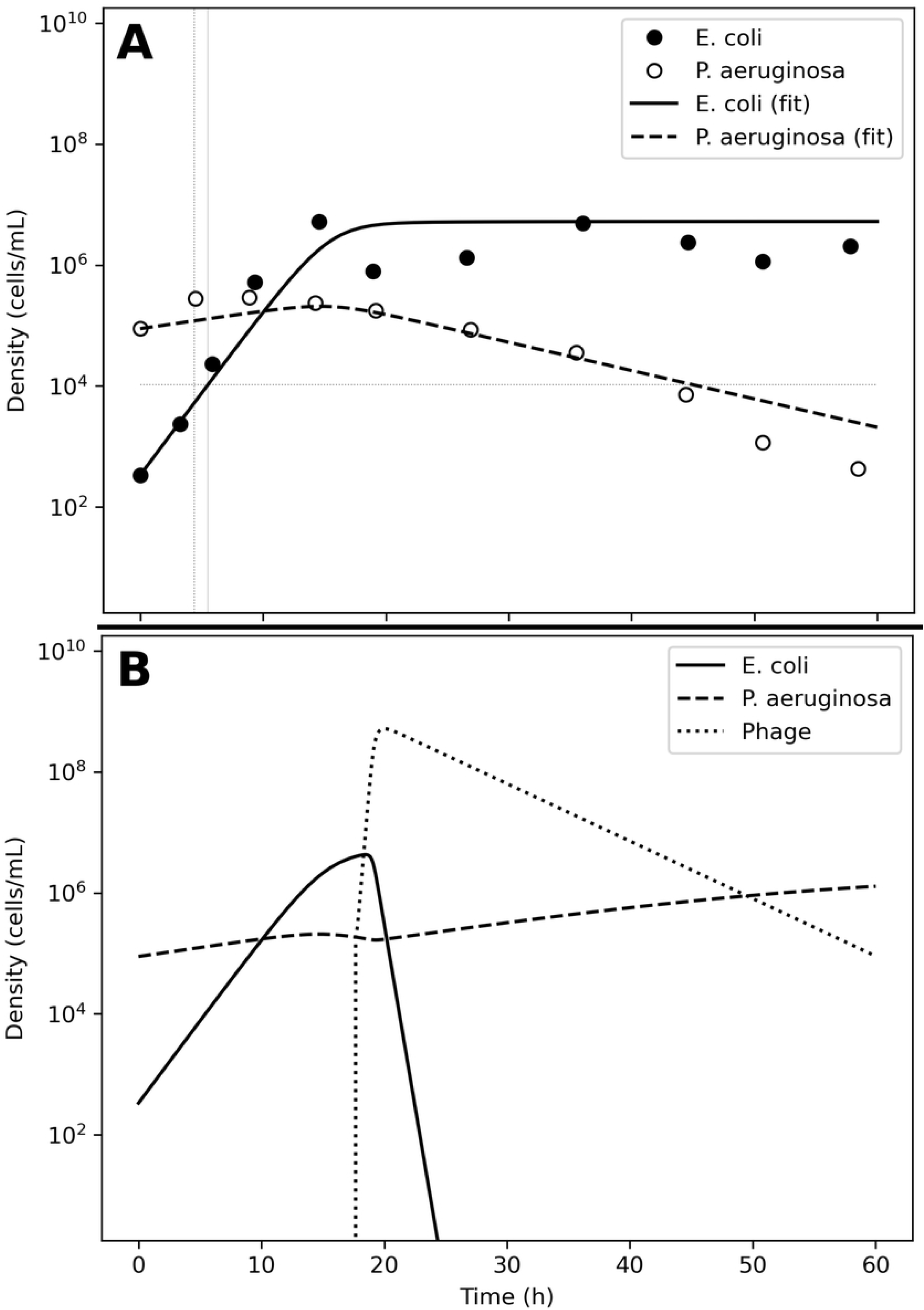
Model of the competition between *Escherichia coli* and *Pseudomonas aeruginosa*. A. Bacterial competition in absence of phages. The data estimated from the original plots for *E. coli* (●) and *P. aeruginosa* (○) is represented together with the fitting obtained using ODE models for *E. coli* (solid line) and *P. aeruginosa* (dashed line). The horizontal line represents *X_P_* and the solid vertical line *T_P_*. To note that the *E. coli* model line did not intersect *X_P_* at *T_P_* but at a later time (doted vertical line). B. Bacterial competition in presence of phages. At *t_ϕ_* = 17.7 h, an inoculum of 2 × 10^5^ phages (dotted line) was added to the simulated consortium, causing the extinction of the invading bacterium *E. coli* (solid line) and the recovery of the resident species *P. aeruginosa* (dashed line).

To simulate the phage therapy, the life-history traits of the coliphage T4 were retrieved from the literature [10]: *δ* = 5.0 × 10^−10^ min^−1^ = 3.0 × 10^−8^ h^−1^; *τ* = 23 min (resulting in *η* = 2.61 h^−1^); *λ* = 0.068 plaque forming units per hour (PFU h^−1^); *β* = 150 PFU; *ω* = 0.15 mL/h. The simulation time-frame was 60 h. The decision tree identified three possible effective outcomes: passive (accuracy 99.3%), active (accuracy 98%) and delayed active (accuracy 100%). Passive therapy required a minimum of 4.50 × 10^9^ PFU/mL; administered before 99.4 h. Active therapy required up to 4.50 × 10^8^ PFU/mL amount of phage between 11.5 and 57.1 h. The best pair of viral load and administration time for active therapy were identified in 199,965 PFU/mL and 17.7 h (Fig 3B). The best pair of viral load and administration time for passive therapy were identified in 9.94 × 10^9^ PFU/mL and 4.0 h (data not shown).

Remarkably, an oscillation in population density was serendipitously obtained with a viral load of 10^6^ at 10 h determined a first wave of phage expansion followed by bacterial decrease and a second wave of phage expansion that caused the collapse of the host population (Suppl. Fig 1).

**Suppl. Fig. 1.**
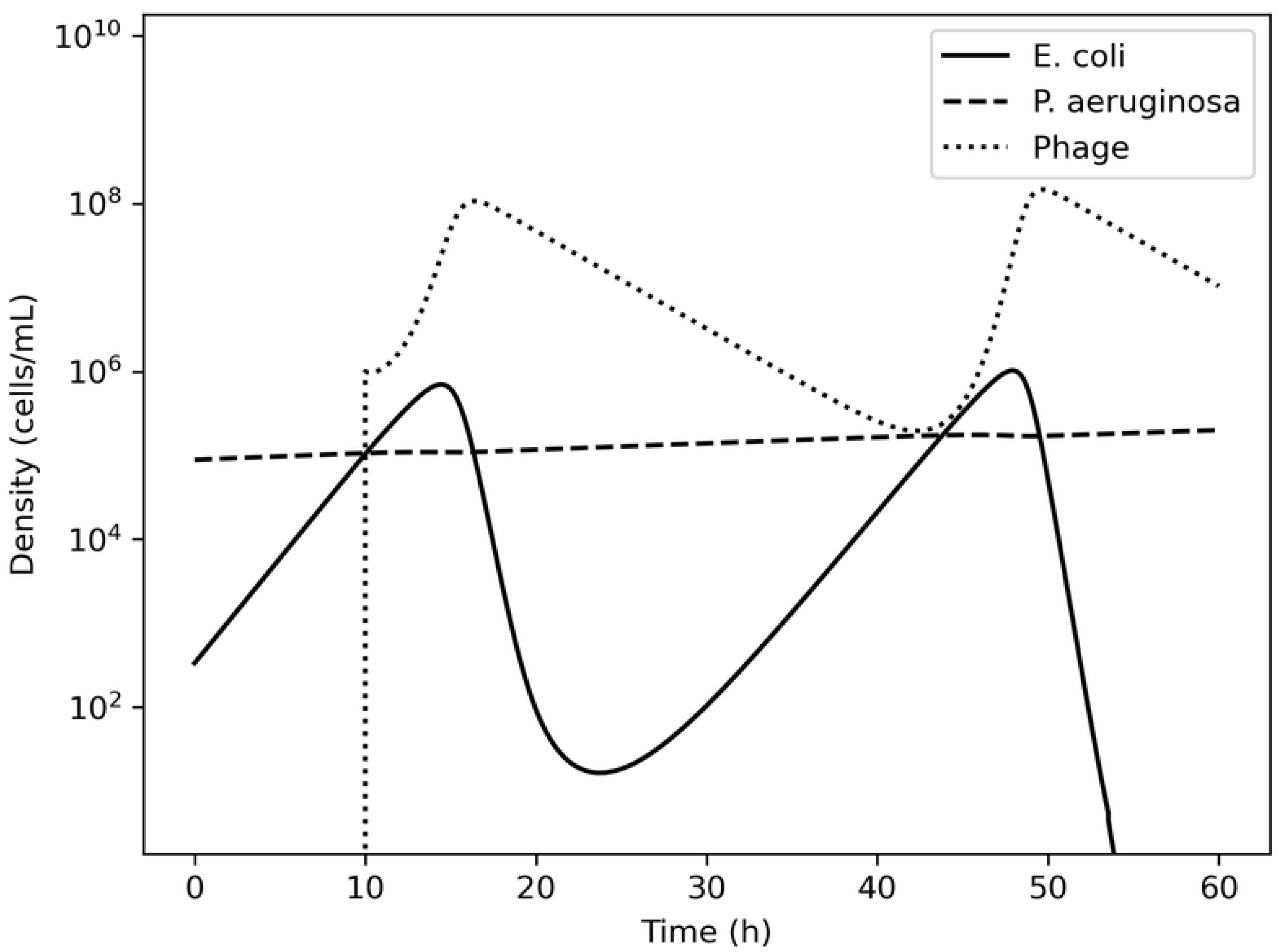
Fluctuation in host-prey density. In certain conditions, bacteria and phages can establish a more dynamic interaction where the depletion of the host is followed by the depletion of the prey, which then results in the expansion of the host. In this case, the second wave of host expansion was followed by eradicating the bacterium resulting in effective treatment. Note how the peak in phagial density precedes the decline of bacterial density.

### Case 3: Escherichia coli vs Azotobacter vinelandii

The growth of the bacteria *E. coli* B/r and *Azotobacter vinelandii* OP was described by Jost and collaborators in 1973 using continuous culture [11]. The authors reported specific growth rates of 0.320 h^−1^ and 0.230 h^−1^ for *E. coli* and *A. vinelandii*, with *K_S_* of 1.0 × 10^−7^ and 1.2 × 10^−2^, respectively. The concentration of glucose in the reactor was 0.005 mg/mL, providing maximum growth rates of 0.320 h^−1^ and 0.070 h^−1^ for *E. coli* and *A. vinelandii*. The calculated growth rate of *A. vinelandii* matched what reported in the public domain [12] but did not allow the building of a fitting model (Suppl. Fig 2). A value of *ν* = 0.200 ± 0.010 was reported in the literature [13] and allowed for a better description of the data (Fig 4A). The data for the simulation were extracted from the original figure of Jost *et al*., providing *X_0_* of 80,251,179 CFU/mL for *E. coli* and 143,462,884 CFU/mL for *A. vinelandii. X_P_* was calculated at 13,258 CFU/mL but, since it was below *X_0_, T_P_* was negative (−27.21 h).

**Fig. 4.**
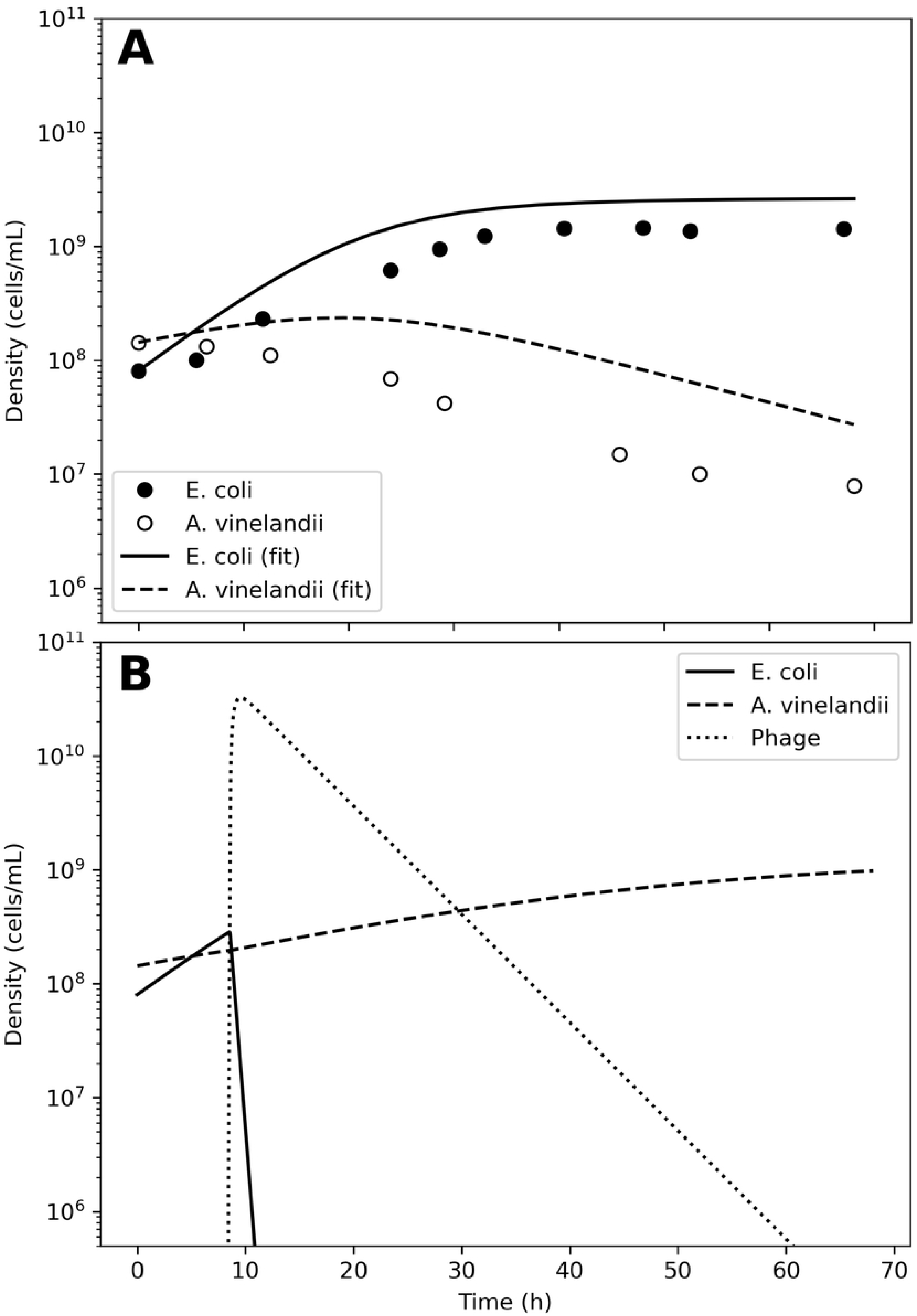
Model of the competition between *Escherichia coli* and *Azotobacter vinelandii*. A. Bacterial competition in absence of phages. The dots represent the data estimated from the original plots for *E. coli* (●) and *A. vinelandii* (○), the lines the conversion to a logistic model. B. Bacterial competition in presence of phages. At *t_ϕ_* = 8.4h, an inoculum of 1.6 × 10^6^ phages (dotted line) was added to the simulated consortium, causing the extinction of the invading bacterium *E. coli* (solid line) and the recovery of the resident species *A. vinelandii* (dashed line).

**Suppl. Fig. 2.**
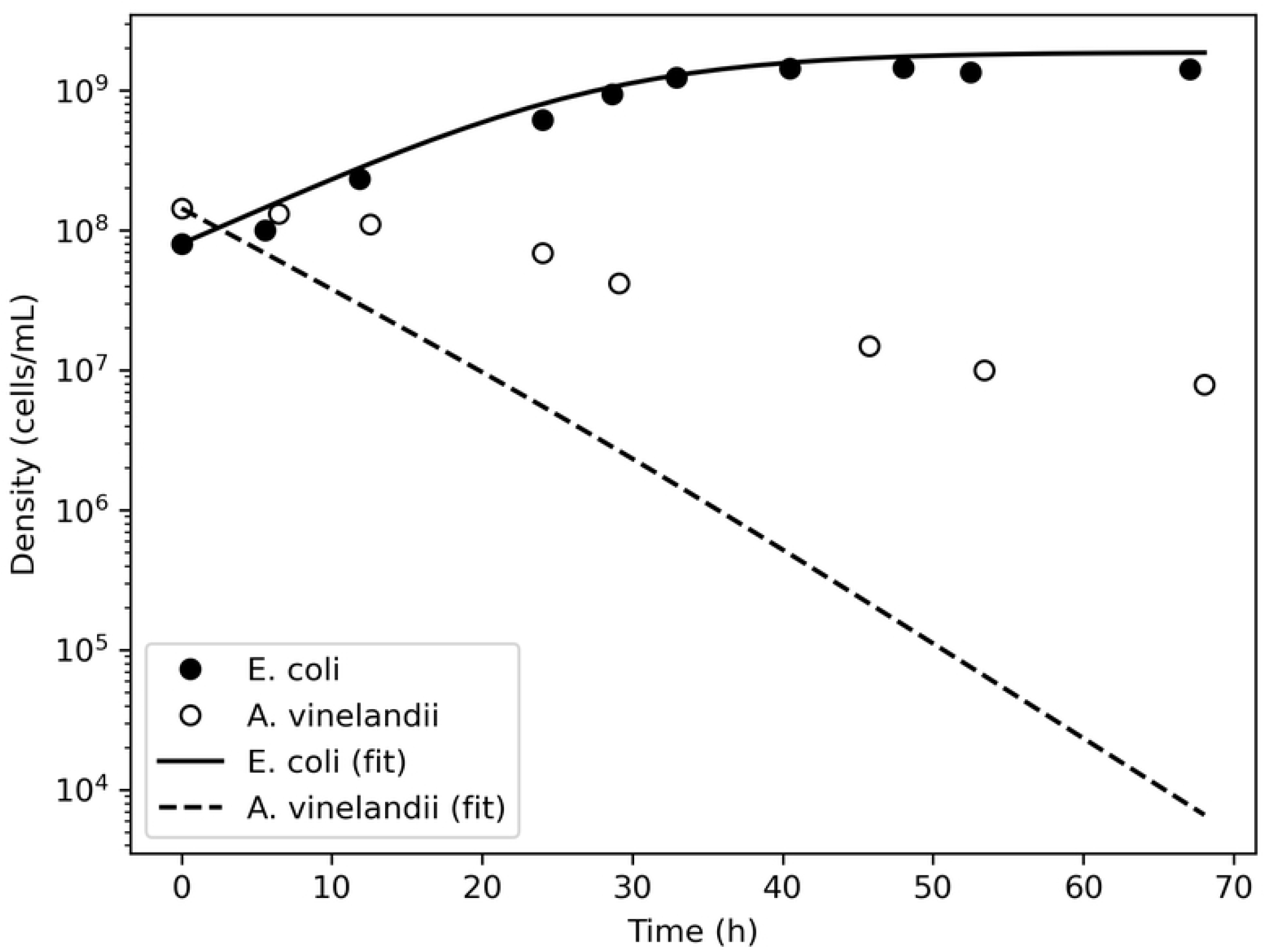
Model of the competition between *E. coli* and *A. vinelandii* using the original growth rates. Plot of the data extrapolated from the study by Jost *et al*. (dots) [11]. The logistic model obtained by using the growth rates reported in the study, 0.32 h^−1^ for *E. coli* and 0.07 h^−1^ for *A. vinelandii*, did not fit the data (lines).

The phage therapy was assumed to use coliphage T4; thus, the life traits were the same as in case 2. The simulation time-frame was 67 h with *ω* = 0.15 mL/h. The decision tree identified two effective therapeutic outcomes: passive (accuracy 100%) and active (accuracy 100%). Passive therapy required at least 1.79 × 10^11^ PFU/mL; any time. Active therapy is obtain below 8.97 × 10^9^ PFU/mL before 66.8 h. The Pareto optimal pair of viral load and administration time for active therapy were identified in 1.63 × 10^6^ PFU/mL and 8.4 h (Fig 4B).

### Case 4: Candida albicans, Streptococcus mutans, and Lactobacillus reuteri

The present case investigated the effect of phage therapy on mutually synergic microbial species. It has been demonstrated experimentally that certain microorganisms inhibit the growth of other microbial species. For instance, *Lactobacillus crispatus* slows the growth rate of both *Gardnerella viginalis* and *Neisseria gonorrhoeae* [14], and *L. brevis* inhibits *Chlamydia trachomatis* [15]. The opposite occurrence is also possible, with microorganisms experiencing increased growth rates when co-cultured with boosting species. For instance, the oral pathogen *Aggregatibacter actinomycetemcomitans* induces higher biomass of *Streptococcus mutans*, a bacterium ubiquitous in the oral flora [16]. Similarly, the pathogenic protist *Candida albicans* also increased the growth rate of *S. mutans* [17]. Microbes can, therefore, influence each other fitness and, consequently, their growth rates.

The case might arise of a phage resistant microbe whose booster species is sensible to phage infection. In that case, targeting the booster species might reduce the virulence of the pathogen and help the clearance of the infection. However, experimental descriptions of such a scenario are virtually absent. Thus, as a proof-of-concept, we defined a hypothetical microbial consortium composed of a phage-resistant pathogen (*C. albicans*, a protist), a boosting bacterium susceptible to phage infection (*S. mutans*), and a commensal bacterium (*L. reuteri*).

The details of the simulation were as follows. Even if not a bacterium, the growth of *C. albicans* has been modeled using logistic models [18], indicating that Eq. 9 (see the *materials and methods* section) was still valid. The growth rates of *C. albicans* and *S. mutans* were estimated from the literature by estimating the cell counts from the original figures [16,17] (Suppl. Fig 3). The density of *S. mutans* in the initial phases of growth in the presence of *C. albicans* was 8.39 ± 6.23 × 10^7^ CFU/mL; conversely, the mean density of *C. albicans* in the presence of *S. mutans* was 1.88 ± 1.11 × 10^6^ CFU/mL. Thus, the ratio *S. mutans/C. albicans* was 44.6. However, these measurements were taken from two different series of experiments, making it difficult to determine an accurate value of *μ_o_* for a single consortium. The growth rate of *S. mutans* was computed at 0.226 h^−1^ when cultivated alone, and at 0.488 h^−1^ when cultivated together with *C. albicans*. Conversely, the growth rate of *C. albicans* was computed at 0.236 h^−1^ when alone and 0.419 h^−1^ when in presence of *S. mutans*. The *L. reuteri* growth rate was derived from the public domain: 0.177 h^−1^ [19] and was considered constant. The model considered an initial seed of 1 × 10^4^ CFU/mL for both *S. mutans* and *C. albicans*, and 1 × 10^8^ CFU/mL for *L. reuteri*. The model showed that both *S. mutans* and *C. albicans* grew with similar dynamics and overgrew *L. reuteri* within 60 hours after the beginning of the simulation (Fig 5A). Specifically, at the end of the simulation, *C. albicans* and *L. reuteri* had densities of 1.2 × 10^9^ and 3.5 × 10^7^ CFU/mL, respectively. Since the growth of the host did not follow a constant trajectory, Eq. 1 was not deemed fitting and no *X_P_* was calculated.

**Fig. 5.**
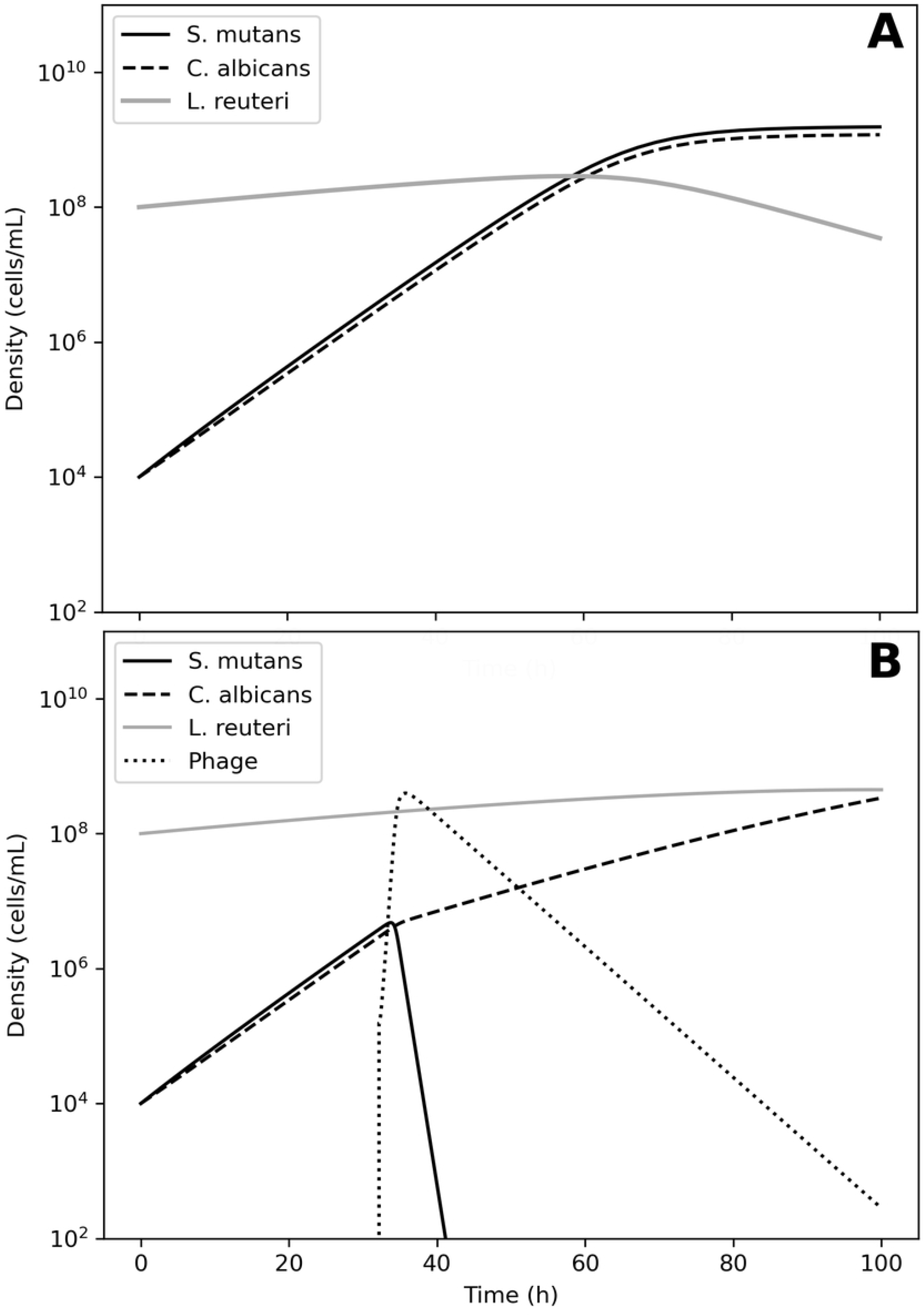
Model of the competition between *Streptococcus mutans, Candida albicans* and *Lactobacillum reuteri*. A. Bacterial competition in absence of phages. Models generated for a hypothetical consortium of two bacteria (*S. mutans* and *L. reuteri*) and one protist (*C. albicans*). The opportunistic species *S. mutans* (solid line) and the pathogen *C. albicans* (dashed line) increase each other growth rates causing a depletion in the commensal *L. reuteri* (gray line). B. Bacterial competition in presence of phages. At *t_ϕ_* = 32.2 h, an inoculum of 1.6 × 10^5^ *λ* phages (dotted line) was added to the simulated consortium, causing the extinction of the *S. mutans* (solid line), a reduction in the density of *C. albicans* (dashed line), and an increase in the density of *L. reuteri* (solid gray line).

**Suppl. Fig. 3.**
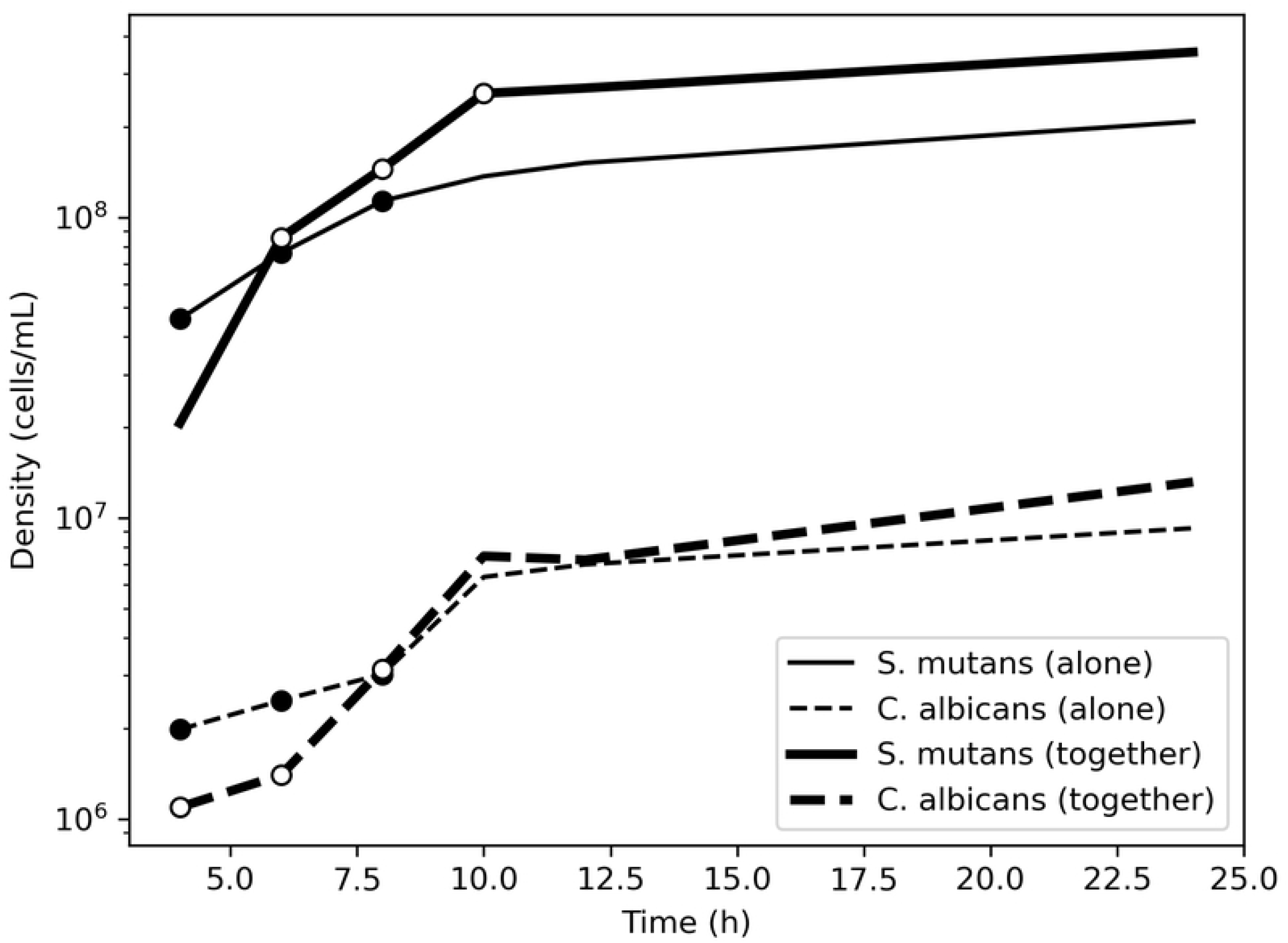
Estimation of growth rates for *Streptococcus mutans* and *Candida albicans*. Densities over time of *S. mutans* and *C. albicans* extracted from the work of Szafrański *et al*. and Sztajer *et al*. [16,17]. The microbes were grown alone or in combination. The data-points used to build the linear models that provided the growth rates are depicted.

The Virus-Host Database reported three phages for *S. mutans:* Streptococcus phage φAPCM01, M102, and M102AD. These phages, all belonging to the family *Siphoviridae*, were highly genetically related: M102 and M102AD shared about 91% similarity at the nucleotide level [20], and φAPCM01 shared 85% nucleotide identity with them [21]. Apart for the M102AD’s adsorption rate (*δ* = 1.5 × 10^−10^ min^−1^ [20]), no other life traits were available in the public domain. Hence, the parameters for the present simulation were derived from another member of the *Siphoviridae* family: phage *λ* [10]. Thus, *δ* = 4.5 × 10^−10^ min^−1^; *τ* = 42 min; *η* = 1.4 h^−1^; *λ* = 0.072 PFU h^−1^; *β* = 115 PFU; *ω* = 0.2 mL/h^−1^; *κ* = 5 × 10^9^ cells/mL. The simulation time-frame was 100 h.

The decision tree identified two possible therapeutic outcomes: passive (100% accuracy) and active (97.3% accuracy). Passive therapy required at least 1.79 × 10^10^ PFU/mL any time. Active therapy required less than 1.42 × 10^8^ PFU/mL virus between 33.0 and 98.3 h. The Pareto optimal pair of viral load and administration time for active therapy was identified in 155,870 PFU/mL and 32.2 h (Fig 5B). At the end of the simulation, *C. albicans* and *L. reuteri* had densities of 3.3 × 10^8^ and 4.4 × 10^8^ CFU/mL, respectively; the peak in phage density was 3.98 × 10^8^ PFU/mL.

## Discussion

Because of the diffusion of multi-drug resistant bacteria, the use of phages to clear bacterial infections is experiencing a resurgence of interest after a long period of obscurity that followed an initial frenzy shortly after its introduction about a century ago. The waning of phage treatment’s initial bliss is due to several concomitant causes, among which a poor understanding of the biological interactions between phages and their host might have led to ineffective treatments [5]. In particular, phage therapy should consider the minimum host density and the rate of phage removal from the micro-environment, with the parameters *V_F_* and *T_F_* determining the thresholds below which the treatment is ineffective [7]. These parameters’ original definitions are formulated as a non-linear equation with two unknowns (the viral load *v_ϕ_* and the administration time *t_ϕ_*). Thus, the calculation of *V_F_* and *T_F_* is cumbersome and, without further conditions, not unique.

In the present study, a numeric approach was taken to identify equivalents for *V_F_* and *T_F_* and to provide the user with a pair of viral load and administration time values equivalent to *v_ϕ_* and *t_ϕ_*. Such an aim was achieved by defining a decision tree that ran the model describing the interaction between phages and bacterial hosts using an ensemble of viral density and administration times and determining the outcome of the interaction. The outcomes were defined, following previous guidelines [7], as: passive, when the virus did not expand above its initial load; active, if the virus expanded above its initial load; delayed, if the virus showed an active behavior after a temporal delay; failed, if the bacterial host was left in the system at the end of the simulation. Since equal outcomes clustered together in the *v_ϕ_/t_ϕ_* space, it was possible to determine the limits of each outcome, obtaining values equivalent to *V_F_* and *T_F_*.

From within the margins for each outcome, it was possible to draw a circle whose area defined a pair of viral load and administration time equivalent to *v_ϕ_* and *t_ϕ_*, which allowed to obtain the chosen type of therapy. The present study’s focus was on active therapy, but it must be stressed that not all combinations of phages and bacteria return every type of treatment. Specifically, passive therapy was achieved virtually in all instances but not so active and delayed therapies. Such an approach was tested on four examples of allochthonous infections.

The first case was derived from the same work that defined the parameters needed for the effectiveness of phage therapy [7]. In their study, Payne and Jansen described a failed passive therapy with the combination of 10^8^ PFU/mL at 2.5 h, and effective passive therapy with the pair 10^9^ PFU/mL at 2.5 h. The results obtained herein indicated that within a time frame of 20 h only passive therapy could effectively clear the infection, and the obtained margins included the values used by Payne and Jansen to achieve effective passive therapy. Conversely, both active and passive therapies could be carried out in the second and third case, with the former also allowing the establishment of delayed therapy.

The fourth case must be treated separately, for the underlying idea in defining the microbial consortium was that there could be occurrences where the pathogen is not a phage’s host. The present study focused on *C. albicans*, an opportunistic fungus that can cause nosocomial infections in multiple organs and it is associated to increased risk of oncogenesis [22,23]. Being a protist, *C. albicans* is immune to phagial infection. However, experimental evidence reported that this pathogen’s growth rate (and consequently its fitness and capability of causing an infection) is increased by booster bacteria, namely *S. mutans* [17]. In the simulation evaluated in the present study, the administration of phages avoided *L. reuteri* being outgrown by *C. albicans*, as instead was the case in the absence of phages. Since *C. albicans* had, in the present study, a growth rate greater than that of *L. reuteri*, the expansion of the fungus would be inevitable, albeit not accomplished within the simulation’s time-frame. The modeling presented in the present study acted as a proof of principle that targeting the booster species will provide, in theory, a ‘mediated phage therapy’ that could reduce a pathogen’s virulence.

The fact that the growth rate of microbes that mutually influence each other ranges between a minimum and a maximum value depending on the density of another species [16,17] implies that ODEs describing such a consortium cannot use constant growth rate parameters. However, the description of such microbial consortia is scanty. Cross-feeding, the metabolic process by which by-products molecules derived from the biochemistry of one species become nutrients for a second species, was the only mathematical description of microbial consortia with dynamic growth rates that could be retrieved [24]. The logistic growth of a species at a density *X* was defined as a function of its growth rate *r* adjusted for the benefit provided by a booster species at a density *Y* and the logistic term:

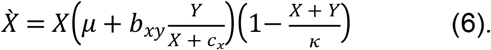

The parameter *b_xy_* indicates the benefit of the species Y over the growth of X, whereas the parameter *c_x_* avoids the problem of division by zero when *X* is equal to zero but does not represent a biological capacity.

An alternative description was presented in the current study. The dynamic nature of the growth rate term in mutual consortia was described by the function *M* (Eq. 13, *materials and methods* section) that shifted the growth rate of the species when grown alone (*μ_ε_*) to that of the growth rate when the booster species was present (*μ_o_*). The function *M* was dependent on the relative densities of the two interacting microbes but, unlike Eq. 6, avoided division by zero adapting the Michaelis-Menten function (*ax*)/(*x*+*b*) [25] in a quorum term 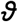, with *a*=1. The property of 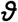 was that it could vary between zero (in the absence of the main species) and one (in the absence of the booster species). The 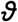 term then decreased *μ_o_* towards *μ_ε_* with decreasing densities of the booster species, dispensing the need for *c_x_*.

The results obtained in the present study confirmed that Eq. 13 was suitable to describe microbial consortia in instances of mutual influence. However, it was impossible to validate the *M* function without experimental data. Alternatives to the Michaelis-Menten function are, in fact, available. For instance, the Holling type III (*ax*^2^(*x*^2^+*b*^2^)^−1^) or the negative exponential (*ae^−xb^*) functions both return unity for *b*=0, with *a*=1, *x*=*ρ*, and 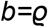. Which of these formulae best describes the growth of mutually influencing microbes could not be determined in the absence of empirical data.

According to a recent review, phage therapy can effectively remove bacteria from a given micro-environment but, in some instances, an invading bacterium can coexist with the resident flora, resulting in a new equilibrium [26]. Should such invading species possess virulence factors, it will be classified as a ‘pathobiont’. It is known that bacteria and phages can establish an equilibrium when having specific life traits and densities [27]. Thus, if the administration of a phage dose does not result in the clearing of its host, it might lead, under peculiar circumstances, to an unforeseen new microbial environment. Suggestion for such an occurrence was observed in the second case, where both bacteria and phages fluctuated prior to the latter’s extinction. Interestingly, the decline in bacterial population followed a peak in phage population, in line with the Lotka-Volterra model [28]. It has been shown that oscillatory conditions between phages and bacteria might occur when the infection rate *η* is within a range whose lower end (*η_c_*) is defined by 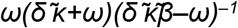, with 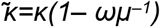 and 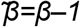 [27]. In case 2, *η_c_* could be calculated in 0.01, which was indeed below the value of 2.61 used in the model.

Discrepancies between the time *T_P_* needed to reach the minimal bacterial density *X_P_* and the simulated bacterial growth were observed. Such a contrast might be due to the fact that the composition of the microbial consortia described herein might not have been suitable for active therapy. Specifically, in the first case, the margins obtained by the decision tree indicated that only passive therapy could be achieved, a treatment that does not depend on *X_P_*, whereas, in the second case, the difference between *T_P_* and the simulated growth was minimal. In both the third and fourth cases, no *X_P_* could be calculated: in the former because the initial host density was above *X_P_*, and in the latter, the dynamic growth rate did not allow for the application of Eq. 1 and Eq. 2. Such a variety of results highlighted the complexity of phage therapy, where each microbial consortium must be considered a unique occurrence whose investigation needs tailored solutions. Overall, the method described in the present work dispensed the need for both *X_P_* and *T_P_*, simplifying the procedure.

The present study had some limitations. The results presented herein were only theoretical and the absence of experimental data strongly limited the validation of the procedures introduced herein. In particular, it was not possible to confirm the validity of Eq. 13 in describing mutually interacting consortia. Another shortcoming of Eq. 13 was that the maximum growth rate *μ_o_* was set in the presence of equal densities of the two interacting species. In reality, the ratio of *S. mutans* over *C. albicans* when determining their *μ_o_* was 44.6 to 1; thus, the quorum term should return one when 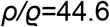. Moreover, Eq. 13 should be generalized to include the cases where the mutual influence between microbes is negative, as in the case of *N. gonorrhoeae* and *L. crispatus* [14]. Again, in the absence of experimental data, the present study was not conceived to achieve such a generalization and additional work is required to better describe the dynamic of mutually influencing species and increase the effectiveness of phage therapy. Another major limitation of the present study was the paucity of life-history traits. The modeling of the microbial interactions had to rely on the scanty data available from the literature or dedicated databases, introducing a bias in the computations. For instance, since the only value available for *S. mutans* phage M102AD was δ, the modeling of case 4 had to be built on phage *λ*, which does not infect this bacterium. In addition, the growth of *C. albicans* was modeled as that of a bacterium [18], an assumption that might not fit the actual growth of this protist [29]. Since, as the present study highlighted, phage therapy needs to be tailored for each microbial interaction, there is a demand to increase the availability of the pool of life-history traits available to microbiologists, a goal that only a multi-center effort could achieve.

In conclusion, the present study introduced machine learning, in the form of a decision tree algorithm, to determine ranges for the phagial dose and administration times needed to achieve passive, active, or delayed treatment. A multi-criteria optimization problem provides Pareto optimal treatment parameters. The procedure used herein dispensed the need for non-linear equations with two unknowns, simplifying the workflow to achieve effective phage therapy. The present study also introduced the concept of ‘mediated phage therapy’, where targeting a booster bacteria might decrease the virulence of a pathogen immune to phagial infection. Such a model required the definition of dynamic growth rates achieved herein with the introduction of quorum terms based on the relative densities of the interacting species.

## Materials and methods

### Microbial growth models

Bacterial growth was implemented using logistic functions and the phage expansion was linked to the bacterial host by the following ordinary differential equations (ODEs):

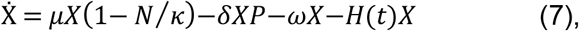

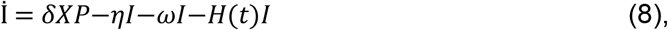

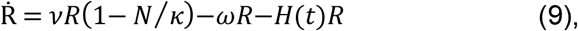

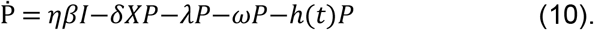

*X* and *I* indicate the population of susceptible and infected bacteria, respectively, whereas *R* is the population of bacteria resistant to phage (*P*) infection. The logistic terms were expressed as the ratio of the total bacterial population *N* to the carrying capacity *κ*. To note that the logistic terms *μX*(1 – *N/κ*) and *νR*(1 – *N/κ*) could be reduced to *μX* or *νR* for *κ* reaching infinity. The use of the variables’ names for the phagial life-history traits followed the suggestions provided by Garcia and co-workers [30]: *β*, burst size; *δ*, adsorption rate; and *λ*, decay rate. In addition, *η* represented the reciprocal of the latency time *τ*, and *ω* the wash-out (outflow) of the microbes. The terms *μ* and *ν* indicated the growth rate of the susceptible/infected and resistant bacteria. *H* and *h* represent the immune response to bacteria and phages, respectively, and are expressed as function of the time of onset and microbial density. Since the examples treated in the present work were derived from *in vitro* cultures, the immune response was ignored; thus, both *H* and *h* were set to zero. The immune response *in vivo* can also be omitted if the treatment is administered quickly enough, further strengthening the assumption of setting both *H* and *h* to zero.

The ODEs’ solver was modified to allow adding the phage dose (*v_ϕ_*) at a given time (*t_ϕ_*). In addition, to avoid a possible reprise of microbial populations that reached very small values (but not zero), an ‘extinction clause’ was added to the solver forcing either *X* or *R* to zero when the associated populations reached fractional values (that is, when the population accounted for less than one bacterial cell). Regression by cubic spline function was used to compute the intersection between the growth models and *X_P_* values [31].

The focus of the present analysis was on what can be described as ‘allochthonous invasion’, that is the outgrow of a species at high cell density by another species initially present at low concentration. This description was based on the definition of autochthonous and allochthonous microflora [32]. An *autochthonous* species is a permanent component of a specific micro-environment, whereas an *allochthonous* species is introduced anew into such a niche. Consequently, the density of autochthonous species is expected to be higher than that of the allochthonous species, but, as in the case of infections, the invading species will eventually outgrow the resident species. Typically, in the context of infection, the autochthonous species can be considered commensal (non-pathogenic to a mammal host, for instance). In contrast, the allochthonous species is usually a pathogen, introduced by several routes such as ingestion or abrasion, due to virulence factors that increase its growth rate over that of the autochthonous species. However, due to a lack of experimental microbial systems, some of the host species included in the cases investigated herein were pathogenic.

The examples used in the present work were derived either from batch (closed vessel) or continuous (chemostat) culture. In the former case, the growth was converted from an explicit consumption of a limiting nutritive resource to implicit consumption under the assumption that the limiting resource would have remained constant. In particular, the specific growth rates were calculated from the maximum growth rates using the Monod term:

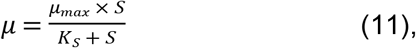

with *S* being the concentration of the limiting nutrient, and *K_S_* being the half-saturation constant [33]. The micro-environment envisioned in the present work was the gut, thus requiring a wash-out parameter *ω* that was set to 0.15 h^−1^ throughout the present work.

### Estimation of growth rates

The microbes’ life traits were based on information retrieved from the literature. When not provided by the experimental settings of the studies considered herein, the growth rates were calculated as a function of the bacterial population at time *t_0_* (*N_0_*) and at time *t* (*N_t_*) with the formula [34]:

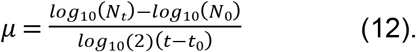

The growth rate was numerically computed as the slope of a linear model based on the bacterial densities displaying a linear distribution.

To account for the shift in growth rates associated to the relative bacterial densities, the ODEs were modified as follows. The growth rate of a given bacterial species alone was indicated with *μ_ε_* (from the Greek ἐρημία: erēmíā, *loneliness*), whereas the growth rate in presence of a booster was indicated with *μ_o_* (from the Greek ὁμαρτῆ: homarte, *at the same time and place*). Since the species started mixing together, the baseline growth rate was *μ_o_*, but a loneliness term *ε* was added to shift *μ_o_* towards *μ_ε_* with decreasing amounts of the booster species. The loneliness term was defined as: 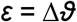, with Δ = (*μ_o_* – *μ_ε_*). The 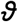 was a ‘quorum term’ derived from the Michaelis-Menten function and was set as: 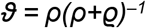, with *ρ* being the density of the influenced (main) species, and 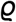 the density of the booster species. The property of *ε* was that it ranged between Δ in absence of booster species 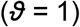 and Δ/*2* when the bacterial densities were equal 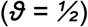. Thus, the constant growth rate of a species (*μ* or *ν*) in Eq. 7 and Eq. 9 was substituted by the function *?* defined as:

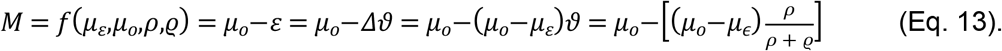

### Ensemble simulations

The analysis of the ODE system (Eq. 7-10) is in general difficult due to the nonlinearity. To study how the viral amount (*v_ϕ_*) and the administration time (*t_ϕ_*) impacted the treatment outcome, an ensemble simulation with 20,000 samples was performed. For each sample simulation, the viral amounts and administration times varied. The values for viral density were randomly selected between 10^2^ to 10^12^ plaque forming units, PFU/mL, with logarithmic scaling. The administration times were equidistant from zero hours to the end of the simulation’s time frame. The range for the viral amount was chosen on the assumption that, while it is possible to make virus dilutions at any desired concentration, administering less than 100 particles per milliliter would have been both impractical and ineffective. Overly concentrated viral suspensions, on the other hand, could produce virion aggregation, reducing the efficiency of the preparation. A topic review of the literature has shown that virtually all phage therapies administer between 10^4^ and 10^9^ PFU/mL; thus, the range provided herein was deemed broad enough to cover all phage therapy situations.

For each of the 20,000 repetitions of the ensemble simulation, the trajectory of the phage was analyzed to determine the type of therapy, classifying it as either active, delayed active, passive or failed therapy. Host concentration above zero at the end of the simulation marked a failed treatment. If the phage concentration never exceeded 105% of the injected amount the therapy was passive. If the phage amount initial decreased before increasing to at least 105% of the injected value, then the therapy was delayed. Finally, if the phage amount directly increased to a value above 105% then the therapy was active.

The generated data of viral amount and administration times (features) and therapy outcome (labels) was used for the decision tree algorithm and to compute Pareto optimal therapy pairs.

### Decision tree algorithm

To compute ranges of viral amounts and administration times for each type of therapy, a decision tree algorithm [35,36] was applied to the output of the ensemble simulation. The decision tree provided a partition of the set of therapy pairs which classified each pair by their expected therapy outcome and the estimated accuracy of the prediction. The resulting ranges gave a simplified representation of the regions of active, delayed and passive therapy outcomes. The boundary of these ranges fulfilled a similar role as the critical values as computed by Payne and Jansen [7]. In comparison to Eq. 4 and Eq. 5, the output of the decision tree did not depend on any asymptotic assumptions on the dynamics of the concentrations. However, they were not as general in the sense that the ranges were only valid for fixed model parameters.

### Pareto optimization

The decision tree-driven classification was not sufficient to select optimal therapy pairs for a specific treatment. For example, therapy pairs at the boundary of the computed ranges are very sensitive to perturbations, resulting in undesirable outcomes for the final user. To compute optimal therapy pairs, we defined a multi-criteria optimization problem (MCOP) [37]. MCOP is widely used to guide the decision of treatment parameters [38]. The criteria employed to achieve an effective therapy was a maximal insensitivity to perturbations combined with the shortest possible administration time. For a given therapeutic pair (*v_ϕ_, t_ϕ_*), the measure of insensitivity was the largest radius *R* of an ellipse such that all perturbed pairs (*v_ϕ_, t_ϕ_*) which satisfied the inequality 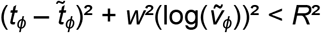 also yielded the desired therapy outcome (Fig 6). The scaling constant *w_ϕ_* determines the shape of the ellipse of perturbations. For all cases in this article, the value *w_ϕ_* = *2* was used. The data from the ensemble simulation provided a fast way to approximate *R*(*v_ϕ_, t_ϕ_*). The MCOP for optimal therapy pairs (*v_ϕ_, t_ϕ_*) was defined by maximizing *R*(*v_ϕ_, t_ϕ_*) and minimizing the time of administration *t_ϕ_*. Such problems could have multiple Pareto optimal solutions, that is, solutions for which all alternatives weaken at least one of the objectives. The weighted sum method [37] in conjuncture with the particle swarm method [39] was used to compute Pareto optimal solutions. Among these Pareto optimal solutions, there was always a unique therapy pair which maximized the insensitivity measure while being reasonably early. The approach used was prototypical in the sense that, depending on the specific application, other criteria could be chosen instead.

**Fig 6.**
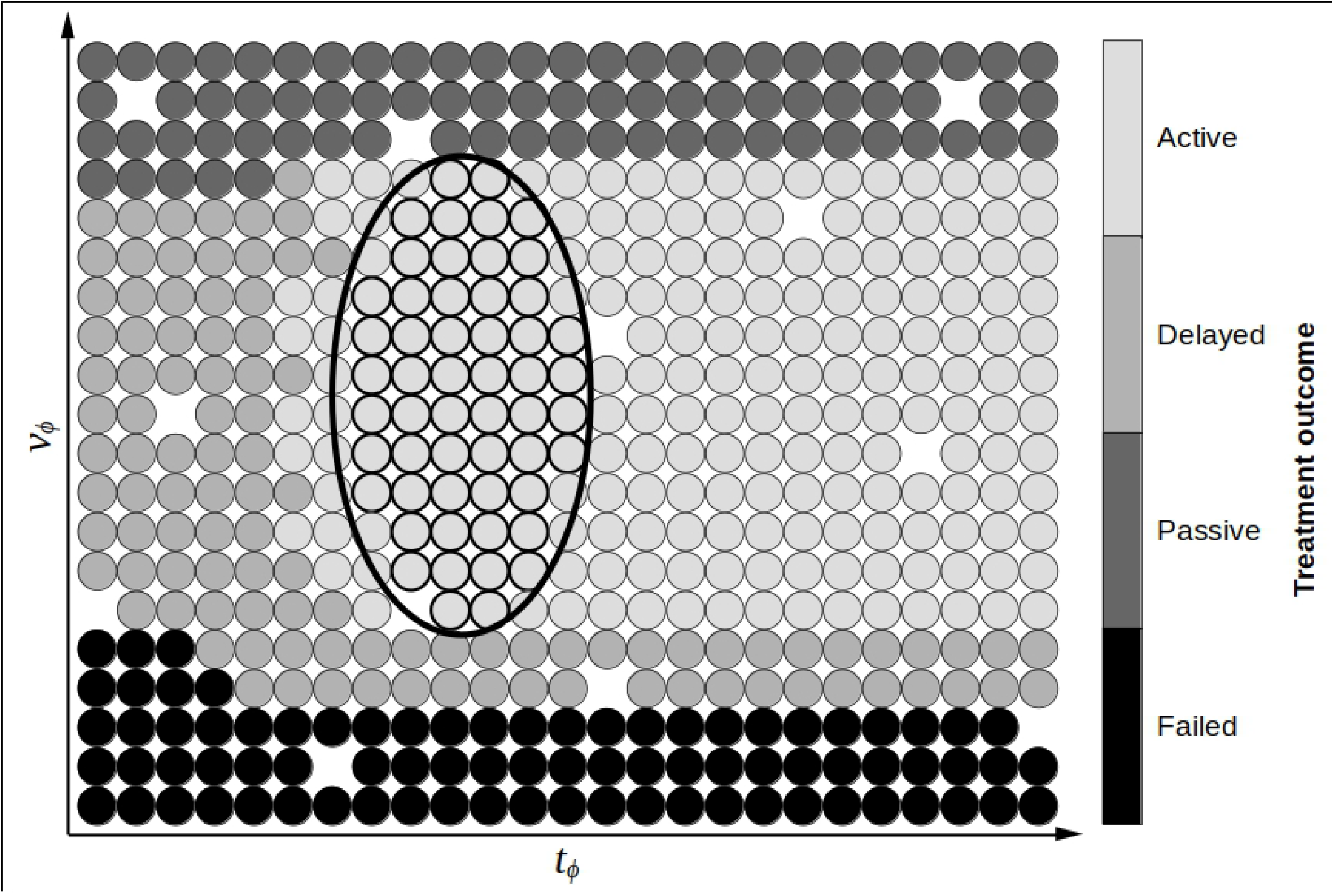
Outlook of the procedure for the selection of the most effective pair of *v_ϕ_* and *t_ϕ_*. The ensemble simulation generates a space of viral dose and administration times whose employment lead to a different outcome, color-coded in the legend on the side of the plot. The boundaries (or margins) between the different regions (spaces of homogeneous color) represent critical values, thus providing the thresholds (equivalent to *V_F_* and *T_F_*) for the range of values required to achieve a specific therapy (passive, active, delayed, or failed). Since the selection was at random, some possible pairs were not computed, leaving the relative positions in the mathematical space undetermined. The selection of optimal pairs of viral load and administration times (equivalent to *v_ϕ_* and *t_ϕ_*) was obtained with a Pareto approach implemented with as a multi-criteria optimization problem (MCOP). This approach is represented by a circle drawn within the chosen therapy (active, in this case) with the characteristics of maximum viral load range and minimal administration time (circle). The center of such a circle provided the user a pair of *v_ϕ_* and *t_ϕ_* that can ease the implementation of the phagial treatment.

### Implementation

Computations were carried out in *Julia* 1.7 [40] and implemented with the packages: *DifferentialEquations* (solution of differential equations) [41]; *LsqFit, Dierckx*, and *Roots* (regression); *DecisionTrees* (classification); *Optim* (optimization) [39]; and *PyPlot* (plotting). Data estimation from the original plots was obtained using *WebPlotDigitizer* 4.5 (https://automeris.io/WebPlotDigitizer/). Bacterial growth rates were computed using a custom function *growthRate*, written in *R* language, that selected the points of bacterial density over time most describing a continuous line and then generated a linear model on those points. The slope of the model was used as the growth rate value according to Eq. 12. Retrieval of phages species for a given bacterium was obtained by inquiring the *Virus-Host Database* [42].

## Acknowledgments

We would like to thank Dr. Szymon P. Szafranski (Hannover Medical School), Dr. Claudia Igler (Swiss Federal Institute of Technology), and Prof. Stephen Abedon (Ohio State University), for their insights on microbial biology. We are also grateful to Dr. Markus Burkard and Dr. Christian Leischner, University of Hohenheim, for critical reading of the manuscript.

## Supplementary data

A dynamic plot of case 1 is available at: https://observablehq.com/@3784219e03ed337e/interactive-phage-simulation.

